# Optic fissure margin morphogenesis sets the stage for consecutive optic fissure fusion, pioneered by a distinct subset of margin cells using a hyaloid vessel as scaffold

**DOI:** 10.1101/141275

**Authors:** Priska Eckert, Lucas Schütz, Joachim Wittbrodt, Stephan Heermann

## Abstract

The optic fissure is a transient gap in the developing optic cup of vertebrates. Persisting optic fissures, coloboma, are a frequent reason for blindness in children. Although many genes have been linked to coloboma, it has remained unclear how the two bi-layered epithelia comprising the optic fissure margins are fusing to form a continuous neuroretina and retinal pigmented epithelium (RPE) respectively. Besides, highly variable morphologies of coloboma phenotypes strongly argue for a diverse set of underlying pathomechanisms.

Here we investigated the contribution of the individual cell types with 4D *in vivo* time-lapse analyses using zebrafish (*Danio rerio*). This allowed defining the respective roles of the participating tissues and cell populations and their activities during fissure morphogenesis, contact formation between the margins as well as during fusion.

We show that optic fissure closure is initiated by a bilateral tissue flow partially in continuation of the dynamic optic cup morphogenesis but additionally including a tissue flow from the optic stalk. This process is followed by the setup of specific fissure margins by a distinct cell population translocating from of the optic stalk. The morphological fusion is triggered by in an EMT-like disassembly of the fissure margin driven by bi-potential pioneer cells that ultimately take the fate of both, neuroretina and RPE respectively. The consecutive fusion and re-epithelialization transforms the two initially separated epithelial bilayers into the two continuous layers of neuroretina and RPE. The processes described here in detail represents a fundamental mechanism of the seamless connection of adjacent multilayered epithelia and is highly reminiscent of other fusion processes, like palatal shelf fusion with key relevance for development and growth.

## Introduction

The optic fissure is a physiological gap in the developing vertebrate optic cup during embryogenesis (Chow and Lang, 2001, Walls, 1942). The fissure is important during a specific period of development, in which it is used by cells of the periocular mesenchyme (POM) and by embryonic vasculature (hyaloid vessels) to enter the eye. It is essential that the optic fissure is closed as development proceeds. A persisting fissure is termed coloboma and is a major cause for blindness in children (Onwochei, 2000). A plethora of genes have been linked to coloboma formation (Graw, 2003), resulting in a coloboma gene network (Gregory-Evans et al., 2004, 2013), which is constantly growing and also includes many signaling pathways, e.g. Wnt (Westenskow et al., 2009, Bankhead et al., 2015), FGF (Chen et al., 2013, Cai et al., 2013), BMP (Heermann et al., 2015), RA (Matt et al., 2008, Lupo et al., 2011), Hippo (Miesfeld et al., 2015) and Shh (Lee et al., 2008). It is noteworthy that the morphology of coloboma phenotypes resulting from pathologies within these genes and signaling pathways is highly variable. Alterations in some of the signaling pathways (e.g. Wnt, Hippo) result in a vast, extended coloboma phenotype, suggesting an early morphogenetic defect (Bankhead et al., 2015, Miesfeld et al., 2015), whereas alterations in another signaling pathway (FGF) result in a more subtle coloboma phenotype (Chen et al., 2013, Cai et al., 2013), potentially involving the alignment or even the fusion of the optic fissure margins. During optic fissure fusion two epithelia, situated in the fissure margins, must be aligned, must get in touch and must fuse, resulting in two new epithelia, the neuroretina on the inside of the optic cup and the retinal pigmented epithelium (RPE) on the outside. However, to this day neither of these sequential processes is understood. Specifically important to understand the complete process would be to understand the following key points: How are the epithelial margins aligned? Which cells establish the contact? How do they do it?

Eye morphogenesis is a dynamic process, involving intensive tissue rearrangements, (Li et al., 2000, Picker et al., 2009, Kwan et al., 2012). We furthermore showed recently, that a precocious arrest of the bilateral neuroretinal flow during optic cup formation results in an extended coloboma phenotype (Heermann et al., 2015). This is a good example for an extended coloboma phenotype and it furthermore suggests that other extended coloboma phenotypes could have a related pathogenesis. Overall, however, the pathomechanism underlying other extended coloboma is only very little appreciated and so is the physiological morphogenesis of the optic fissure. We hypothesize that the bilateral neuroretinal flow (Heermann et al., 2015) is involved, also in the morphogenesis of the optic fissure and is potentially important for the alignment of the optic fissure margins, setting the stage for the consecutive fissure fusion. Here, we used zebrafish (*Danio rerio*) and *in vivo* time-lapse imaging to address the morphogenesis and the consecutive fusion of the optic fissure margins.

We found that cells are integrated ventrally into the optic cup from the lens-averted domain and the optic stalk. These movements were bilateral, split in a nasal and a temporal domain and using the emerging gap for involution. This gap is the optic fissure and the process observed was its morphogenesis. Notably, we observed that distinct cells were added to the optic fissure margins only very late from the region of the optic stalk. Furthermore, we identified a population of cells, which leave the epithelial margins as pioneers in an epithelial to mesenchymal transition (EMT)-like process. These cells use the hyaloid vessel as a scaffold to bridge the fissure and eventually establish the contact between the margins. These cells showed morphological features of RPE cells, however, our data indicate that they were bi-potential and their progenies participated in the neuroretina and the RPE. Our data indicate that this process occurs regularly in the upper third of the fissure. Since the pioneering cells mark the border/edge between the future neuroretina and RPE the tissue flow, through the optic fissure must invaginate this border into the fissure.

Our findings provide profound insights into the sequential mechanisms, which are necessary for optic fissure fusion. These findings can likely, at least in part, be applied also to other fusion processes e.g. during palatal fusion and lip fusion during oro-facial development or neural tube closure during spinal cord development.

## Results and discussion

### Pioneering edge cells mediate optic fissure fusion

We first addressed the process of optic fissure fusion and investigated the population of cells mediating the contact in between the optic fissure margins, by using in *vivo* time-lapse imaging of zebrafish embryos. We observed that the onset of optic fissure fusion was occurring proximally and that the process was extending distally, in line with recent observations (James et al., 2016). We focused on distal optic cup domains, which are accessible for high quality *in vivo* time-lapse confocal imaging. Confocal stacks were recorded from a lateral perspective (Figure 1A, oriented nasal to the left), providing optimal resolution in x and y axes. We have previously shown that an *rx2* cis regulatory element is well suited to drive expression in all retinal progenitor cells (Heermann et al., 2015). Thus, we made use of the *tg*(*ola.rx2:egfp-caax*) zebrafish line in which the *rx2* cis regulatory element is driving GFPcaax in all retinal progenitor cells. In the prospective RPE cells, however, *rx2* driven expression was ceasing over time (Figure 1 B, C). To introduce also an *rx2* independent label, for the *rx2* negative population of retinal progenitor cells, we made use of RNA injections into zygotes, with RNA coding for lyntdTomato (Figure 1B-J). Addressing optic fissure fusion over time (Figure 1 B-C, supplemental movie file 1, first half, see Figure S1 for single channels), we identified cells, which as pioneers extended their processes through the fissure, eventually mediating the contact in between the optic fissure margins, thereby initiating the fusion. Notably, these cells were positive for lyntdTomato but largely to completely negative for *rx2* driven GFP (Figure 1C, white arrow). The flattened morphology of the pioneering cells in combination with the terminated *rx2* driven expression was indicative for an RPE precursor fate. Thus, it was very surprising that the rx2 driven expression was reactivated after the fusion was achieved. This is essentially what we observed when we followed the domain of the former fissure margins during further development (Figure 1 D-F, see Figure S1 for single channels).

**Figure 1.**
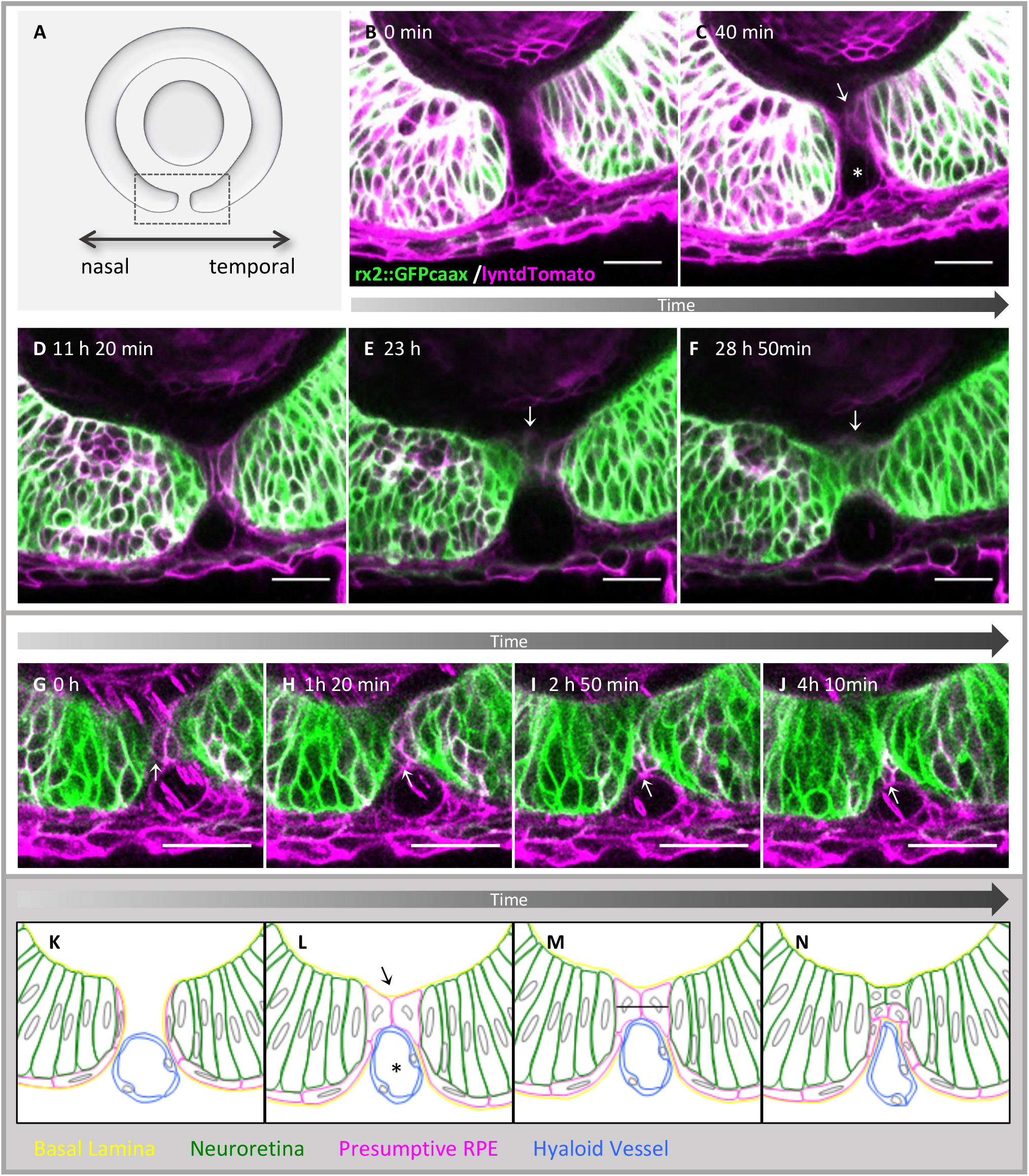
Pioneering edge cells establish the contact of the fissure margins and subsequently become precursors of NR and RPE. (A) Scheme of a lateral view on the developing optic cup for orientation, the boxed region was recorded in B-J. (B) Optic fissure margins before contact formation, *rx2* negative, lyntdTomato positive cells line the nasal and temporal fissure margins. (C) Pioneering edge cells establish the contact between the fissure margins (arrow). (D) Optic fissure after the contact was established. (E) Domain in which the contact was formed reactivates *rx2* driven GFP expression (arrow). (F) Optic fissure after fusion, rx2 is expressed in the contact forming cells (arrow). (G-J) Pioneering edge cell, lyntdTomato positive and *rx2* negative, on the temporal fissure margin establishes the contact between the margins and is subsequently integrated into the RPE (arrow). Scalebar: 25μm, (K-N) Scheme of optic fissure fusion, (K) *rx2* negative cells line the fissure margins (magenta) a basal lamina (yellow) is covering the presumptive RPE and the neuroretina. Between the margins the hyaloid vessel (blue) is visible. (L) Over time, the basal lamina is degraded and the pioneering edge cells (magenta) form contact in between the fissure margins using the blood vessel as scaffold. (M) Pioneering edge cells divide and the progenies are integrated in either the future RPE or neuroretina (N), *rx2* driven expression is reactivated in the progenies which were integrated into the future neuroretina.

These findings indicate, that these pioneering cells are likely not a population of RPE precursors, nor a simple neuroretinal precursor population.

The dynamic expression of the *rx2* cis regulatory domain, observed in the pioneering cells, was reminiscent of the *rx2* expression inside the ciliary marginal zone of the eye, where *rx2* gene expression is active in the stem cells, is lost in the transit amplifying zone and then reactivated in cells of the differentiating retina (Reinhardt et al., 2015). This analogy is especially tempting, since during development, the future CMZ and the optic fissure margins are one continuous structure. Noteworthy, the CMZ contains stem cells for the neuroretina and the RPE, potentially a single population of stem cells.

Hence, we asked next, to what extent also prospective RPE cells are derived from the population of pioneering cells. Notably, in two cases, RPE precursors in the fused domain could clearly be detected as progenies of the pioneering cells (Figure 1 G-J, arrows, supplemental movie 2, see Figure S1 for single channels). In other cases, these analyses were hampered by the pronounced flat morphology of the differentiating RPE precursors, which in combination with limitations in spatial and temporal resolution during *in vivo* time-lapse imaging made cell tracking difficult. Taken together our data indicate, that the pioneering cells are initiating the optic fissure fusion and consecutively differentiate into neuroretina and RPE, suggesting that they are bi-potent progenitors. Considering this, it becomes clear that at least one cell division is required to integrate one descendant into the neuroretina and one descendant into the RPE, which are henceforth located head to head on two separated epithelia (Figure 1K-N, scheme). Notably, defects in cell proliferation have been demonstrated to affect the closure of the optic fissure. Pax2, in combination with Vax2, was shown to be important for cell proliferation and also cell death within the optic cup via regulation of the Fadd gene (Viringipurampeer et al., 2012), suggesting to be the reason for coloboma in the pax2 related renal coloboma syndrome (Torres et al., 1996, Favor et al., 1996, Fletcher et al., 2006, Bower et al., 2011). Independently, abnormal cell cycle progression was also shown to be the reason for coloboma in the Phactr4 KO mouse (Kim et al., 2007). Having shown that the pioneering cells are participating in the formation of the neuroretina and the RPE, it becomes clear that they also mark the border or edge between these precursors of these two epithelia within the optic fissure margins. We thus also named these cells “pioneering edge cells”. We further noticed that at the time, these pioneering edge cells initiated the fusion, the border in between the prospective neuroretina and RPE was inverted into the fissure (Figure 1B-C).

### Hyaloid vessel used as a scaffold by the pioneering edge cells

Besides the pioneering cell population itself, we identified a nearly circular hole over which these cells extended their protrusions and eventually their cell bodies (Figure 1C, asterisk). The morphology and especially the dynamics of fast cell movements inside this structure seen during *in vivo* time-lapse imaging (supplemental movie files 1) suggested, that this structure is corresponding to a blood vessel. To substantiate this finding we performed time-lapse imaging using a transgenic line in which endothelial cells are labelled (*tg*(*fli:GFP*), Lawson und Weinstein, 2002) (Figure 2A-C). Indeed, this structure corresponds to a blood vessel (Figure 2A-C, asterisk, supplemental movie file 3). Taken together, our data indicate, that a part of the hyaloid blood vessel, which is located in between the fissure margins, is used as a scaffold for the pioneering edge cell population to get in touch and to bridge the gap of the fissure (Figure 1C, Figure 2A-C, magenta). We hypothesize that the extracellular matrix (ECM) of the hyaloid vessel segment serves as a grid for the pioneering cells to do so. Recently the hyaloid vessel was suggested to play a role during degradation of the basement membrane of the fissure margins (James et al., 2016), a step which must precede the onset of fissure fusion. A zebrafish mutant lacking hyaloid vessels (cloche, Stainier et al., 1995, Dhakal et al., 2015, James et al., 2016) was used by James and colleagues to show a delayed breakdown of the basement membrane in that situation. Based on our findings we suggest that the hyaloid vessel also plays an important role during fusion itself. An abnormally large vessel due to a *lmo2* mutation was shown to result in coloboma (Weiss et al.,2012). This could mean that the gap, the pioneering edge cells have to bridge, is too big, but it could also mean that in this case the ECM composition was not allowing the pioneering edge cells to use the vessel as a scaffold sufficiently. We noticed that the region of basement membrane degradation indicated by anti-laminin immunohistochemistry (Figure 2D-F) was in line with the region of fusion onset, observed during *in vivo* imaging. To furthermore address the exact location of the fusion area with respect to the total height of the optic fissure we measured the height of the contact points in individual embryos and related these to the total height of the optic fissure (Figure 2G). Our data indicate an onset of optic fissure closure in the upper third of the fissure. This suggests that the border in between the prospective neuroretina and RPE and with it the pioneering edge cells have to be brought into this position, to enable the fusion of the margins. As mentioned above it was suggested that the hyaloid vessel is somehow involved in basement membrane degradation (James et al., 2016) preceding optic fissure fusion. However, it was also discussed that the POM derived precursors of the hyaloid vasculature could actually be the cells, relevant for this. Cells of the POM were previously linked to optic fissure closure, in the context of RA dependency of this process (Lupo et al., 2011). Here, we use *in vivo* time-lapse imaging to investigate cells of the POM in the optic fissure. We make use of a *sox10* cis regulatory element driving a membrane localized GFP (*T2*(*sox10:GFPcaax*)). To visualize individual cells, we performed time-lapse imaging of embryos injected with the construct at the zygotic stage, resulting in a mosaic labeling (Figure 2G-I, supplemental movie file 4). Our data indicate that *sox10* positive cells of the POM are migrating into the optic fissure and, notably, get in touch with the optic fissure margins with long cellular processes. By this contact these cells can potentially induce changes to the ECM within the optic fissure margins and its basement membrane, supporting the idea that POM cells are important for basement membrane dissolution.

**Figure 2.**
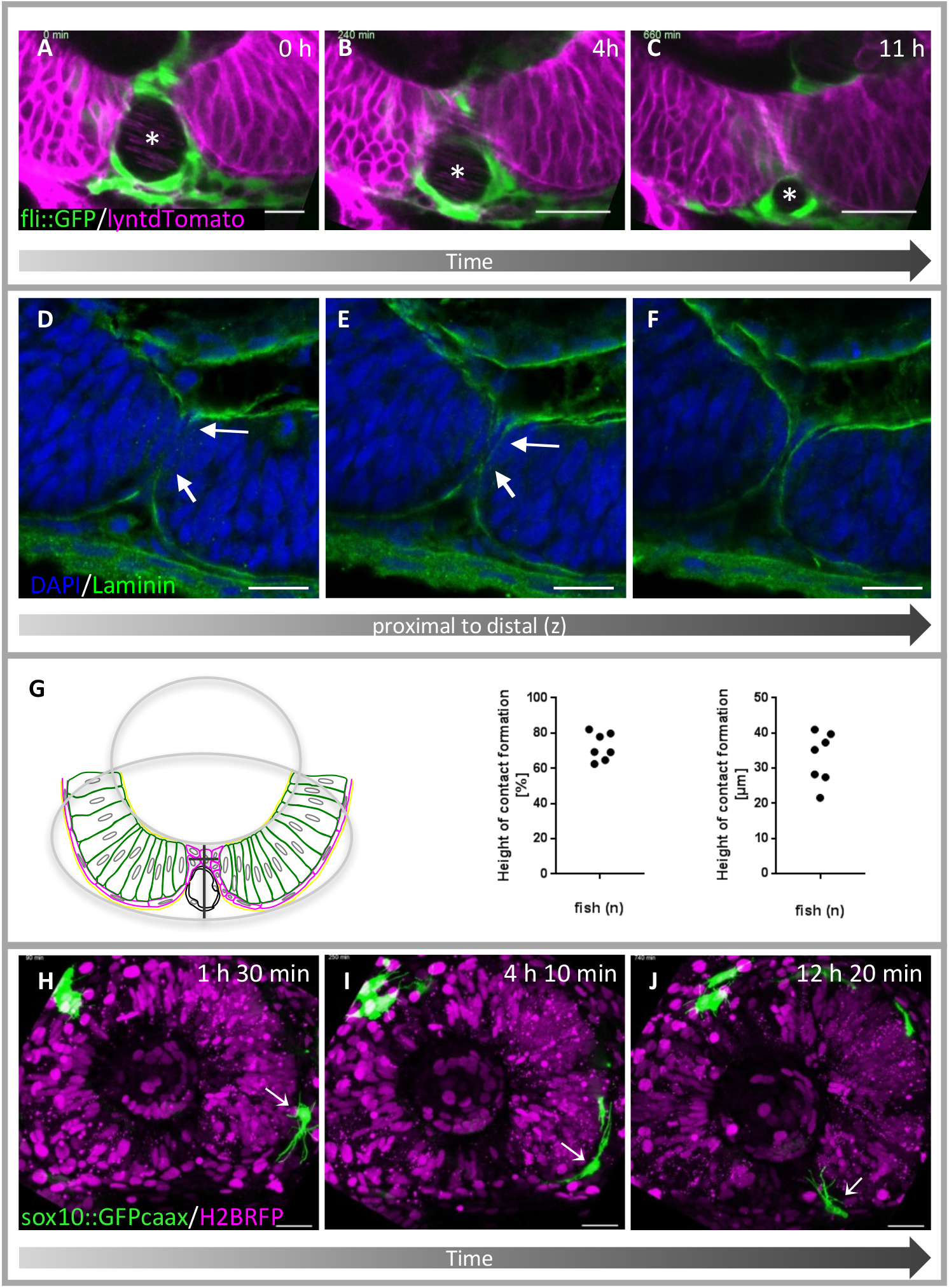
Pioneering edge cells establish contact using the hyaloid vessel as scaffold. (A-C) Pioneering edge cells (magenta) establish contact using the hyaloid vessel (asterisk)(green) as a scaffold, to bridge the gap between the margins. (A) Fissure margins prior to fusion, the hyaloid vessel occupies the room between the margins. (B) Contact formation of pioneering edge cells on top of the vessel. (C) Fissure margins fused over the vessel, which is moved ventrally. (D-F) Degradation of laminin (green, imunohistochemistry) during contact formation from proximal to distal (left to right). (D) Proximal: basal lamina is degraded where fissure margins have contact (between arrows). (E) Degradation of the basal lamina is ongoing between the arrows, to allow the margins to fuse. (F) Distal basal lamina is still intact. Scalebar: 15μm, (G) Measurement of the height of the contact formation between the fissure margins. The height was measured according to the scheme presented. The total fissure height was set as 100%. The pioneering edge cell was virtually split in half and the distance to the respective borders was measured from that point. This was performed for 7 individual embryos. In all cases the height of the fusion point is located in the upper third of the optic fissure (60–80% of the total height). Expressed in total numbers, the contact formation takes places at a height of 20-40μm. (H-J) A periocular mesenchyme (POM) cell (arrows) moving around the optic cup into the optic fissure in a highly motile way, building and retracting protrusions. Scalebar: 25μm

### Optic fissure morphogenesis and the role of the bilateral neuroretinal flow

In the consecutive part of the project we addressed the morphogenesis of the optic fissure and specifically asked how the optic fissure margins are formed during development. With respect to the structure and morphology of the optic fissure margins during fissure closure, two conflicting scenarios were discussed based on long standing observations. On the one hand, it was proposed that the neuroretina would be everted to the outer surface (Mann, 1950, Duke Elder, 1958) on the other hand the RPE should be inverted into the optic fissure (Hero, 1990). A lot of evidence including our own data (this manuscript) is suggesting the latter scenario to be accurate, however, it is intriguing to also consider the conflicting data. Our recent observation that a bilateral neuroretinal flow from the lens-averted into the lens-facing domain of the optic cup is largely contributing to optic cup formation (Heermann et al., 2015) suggested that this bilateral flow is also contributing to the formation of the optic fissure. We now specifically address the neuroretinal flow with respect to optic fissure generation to investigate its contribution to this process. We make use of *in vivo* time-lapse imaging to this end. At first, we performed time-lapse confocal imaging using a ubiquitous labeling for nuclei and membranes (*tg*(*bact2:H2BGFP*, *bact2:lyntdTomato*)) (Figure 3A-F and 3A’-F’, scheme, supplemental movie file 5,6). While the neuroretinal flow over the distal rim is difficult to follow in a lateral perspective, the ventral flow can be nicely appreciated (supplemental movie file 5). Our *in vivo* imaging data clearly show that indeed prospective neuroretinal cells are secondarily added to the most ventral aspect of the optic cup. Importantly, this flow is bilateral, thus splitting the ventral domain of the optic cup, resulting in the formation of the optic fissure (Figure 3A-C, dotted arrow). Notably, cells are added to the temporal-ventral domain mainly from the lens-averted domain of the optic cup, whereas the cells, which were added to the nasal-ventral domain were largely stemming from the optic stalk (Figure 3B, supplemental movie file 5). We furthermore observed, that the connection of the optic stalk to the nasal optic fissure margin was broader than its connection to the temporal margin (Figure 3B). Besides, a vortex of cells was visible in the ventral nasal domain during fissure formation (Figure 3D,E, #, Figure 3S, supplemental movie file 7), which can potentially be explained by cells moving from the lens-averted domain over the optic stalk into the ventral nasal domain.

**Figure 3.**
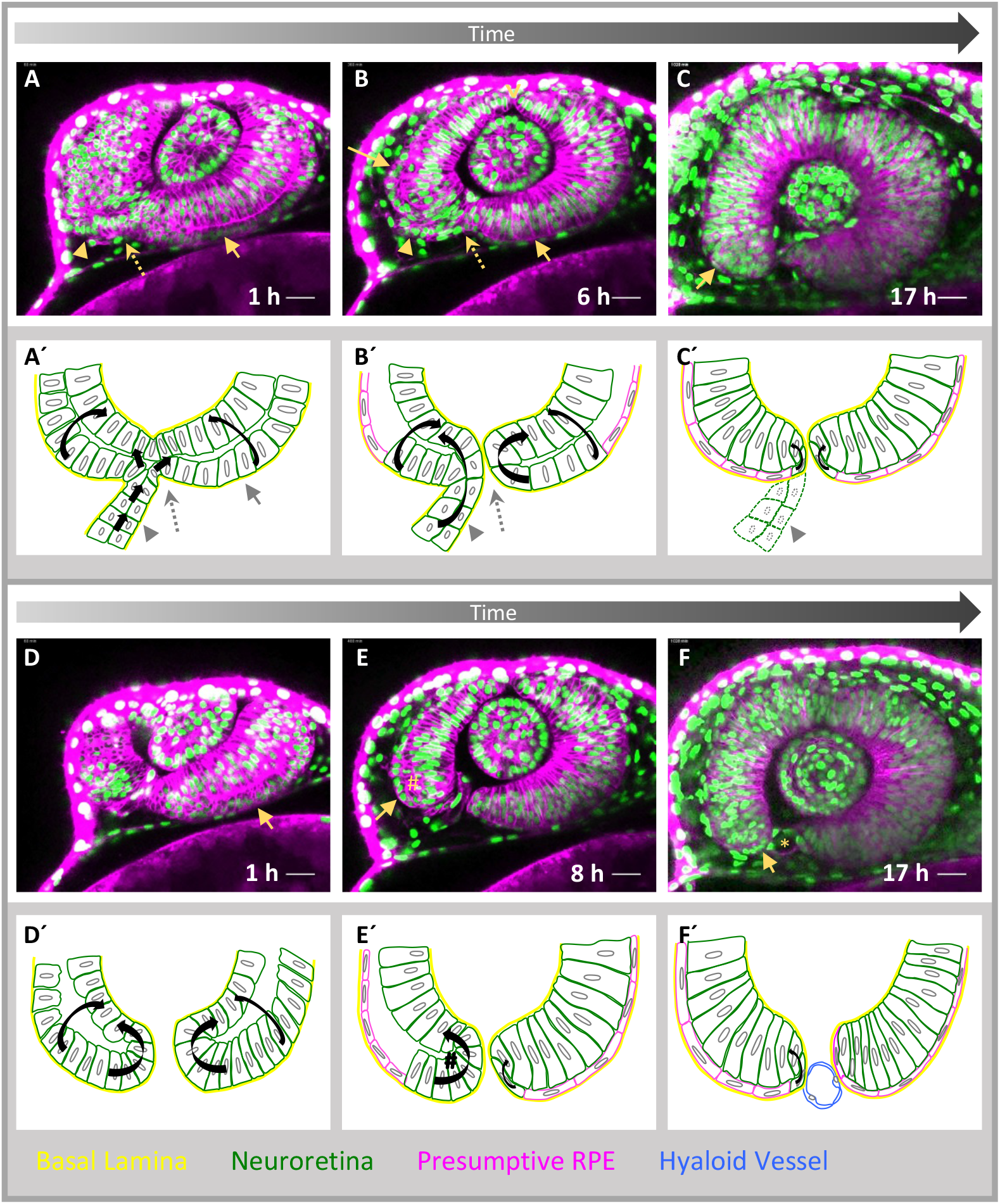
Morphogenesis of the optic fissure (lateral perspective, nasal to the left) (A-C) Proximal region of the developing optic cup over time, (A’-C’) corresponding schemes, (D-F) Distal region of the developing optic cup over time, (D’-F’) corresponding schemes (D+ D′). Cells from the lens-averted optic cup domain (A, A′, B, arrow) and the optic stalk (A, A′, B, B’, arrowhead) move towards the lens-facing domain of the optic cup (A′, B’, black arrows). Moving Cells take two distinct routes, over the distal rim (A′ B’ E’, curved arrows) and notably also bilaterally through the forming optic fissure indicated by a dotted arrow, (A’-F’). Cells remaining in the RPE domain flatten and thus obtain RPE cell shape (B-F, arrow, B’-F’, magenta colored cells). The optic stalk is predominantly connected to the nasal fissure margin (A, A’, B, B’ C’, arrowhead). Cell movements over the distal and ventral rim in this domain (D′, E’, curved arrows), result in a cellular vortex (E, E’, #). The bilateral neuroretinal flow over the distal rims is resulting in a dorsal indentation (B, empty arrowhead) occasionally referred to as dorsal fissure. The hyaloid vessel is visible (F, asterisk, F′, blue), * marks the optic ventricle. Scalebar 25μm

The finding of a bilateral flow, splitting the ventral optic cup, suggests that a domain of the ventral proximal aspect of the optic cup must be immobilized, in order to split the flow of tissue into a nasal and a temporal domain. The future optic nerve head is located in the most proximal domain of the optic fissure and thus likely corresponds to the immobilized domain.

To facilitate a better individual tracking of cells, we reduced the density of labelled nuclei by injecting RNA coding for H2BGFP into one cell of a 4 to 8 cell blastula in the context of a ubiquitous membrane labeling, achieved by injections of RNA coding for lyntdTomato into the zygote. In addition, we wanted to increase the resolution in z during time-lapse imaging. We thus made use of single plane illumination imaging (SPIM lightsheet, Figure 4 A-H, supplemental movie file 8). We tracked single cells on their way into the ventral optic cup. Supporting our data (Figure 3) cells moved on both sides, bilaterally, over the ventral rims of the forming optic fissure. Notably, on the nasal side we found the optic fissure margin connected to the optic stalk and cells moving over the ventral margin in this domain were stemming from the stalk region (Figure 4 A-D). Further supporting this, we found that these cells considerably moved in z-direction. On the temporal side (Figure 4 E-H) we found, next to the cell movements over the ventral rim, also movements over the ventral distal rim and the distal rim. This finding indicates, that the bilateral neuroretinal flow over the distal rim, described previously (Heermann et al., 2015), is in continuation with the flow through the temporal optic fissure margin. In the nasal side, however, the flow is a composite of the neuroretinal flow over the distal rim and the flow, entering from the optic stalk. We hypothesize that turbulence is formed in the border region, in which these two flows overlap, likely resulting in the cellular vortex, which we found in exactly this domain (Figure 3, Figure S3).

**Figure 4.**
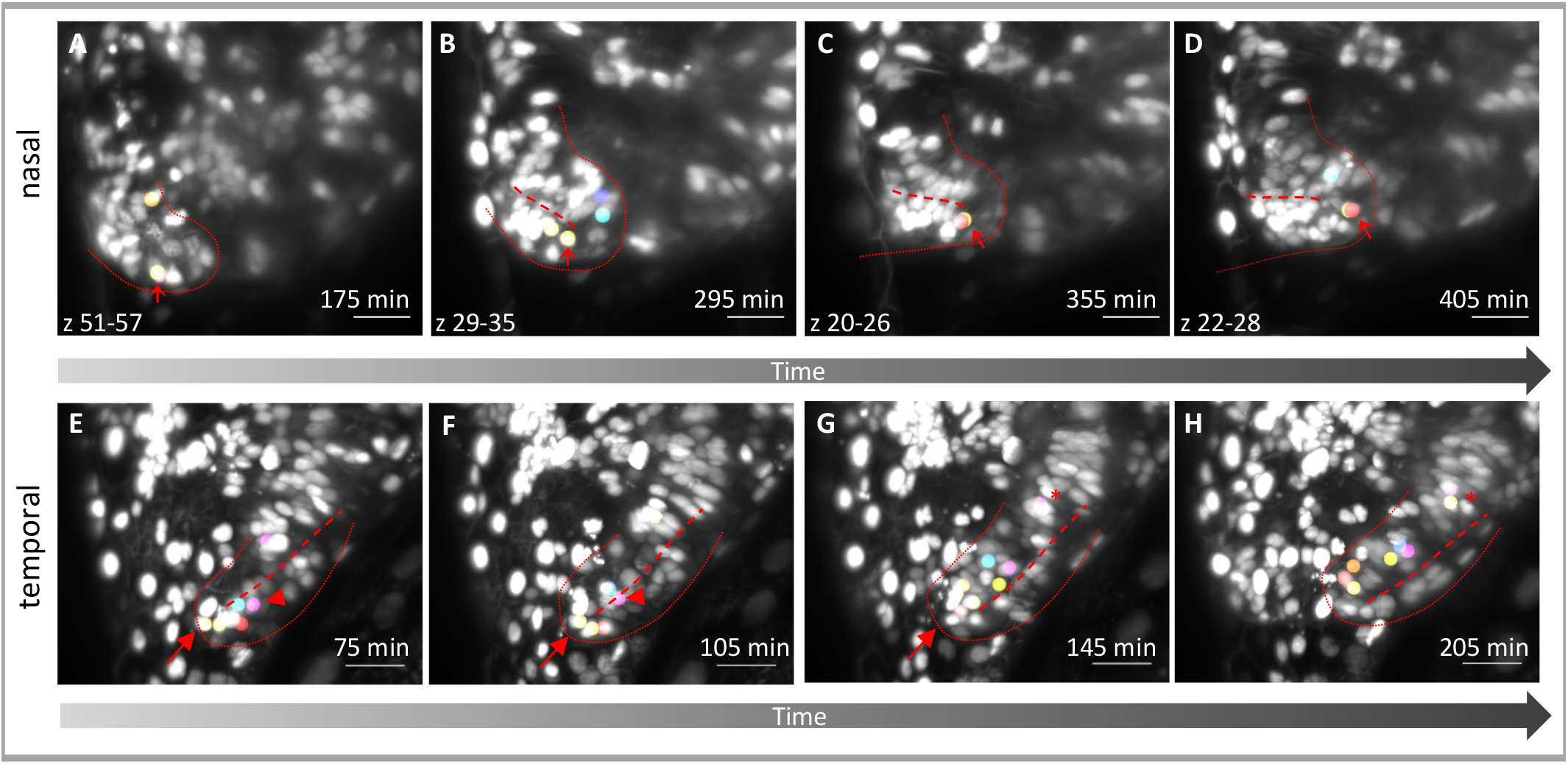
single cell tracking during bilateral ventral flow. A mosaic nuclear labeling facilitated tracking of single cells through the optic fissure over time. Z-projections of the nasal (A-D) and temporal (E-H) optic fissure margin development. Tracked cells (colored dots) can be followed on their way into the optic cup. A dotted curve is indicating the optic fissure margin, which is nasally connecting the optic stalk with the lens-facing optic cup domain (A-D) and temporally connecting the lens-averted domain with the lens-facing domain of the optic cup (E-H). A dashed line (B-H) indicates the border between these domains and marks the ventral rims. Especially on the nasal side, cells moved considerably in the z-plane. In order to still visualize the tracking results, different optical sections had to be used for the z-projections (A-D). Please note the tracked cell (arrow) moving from the optic stalk (A,B) over the ventral rim (C). On the temporal side (E-H) cells can be tracked over the ventral distal rim (E,F, arrowhead, e.g. purple and blue cell) and the ventral rim (E-G, arrow). In addition, in more dorsal domains cells entered over the distal rim (G,H asterisk). Scalebar: 25μm

With respect to the contradictory observations introduced at the beginning of this section (Mann, 1950, Duke Elder, 1958, Hero, 1990), our data suggest, that both observations are accurate. However, it is very likely that the observations resulting in the statement that the neuroretina must be everted for closure were made at a slightly earlier point of time than the observations resulting in the statement that the RPE must be inverted for closure.

### Additional TGFβ signaling positive cells enter from the optic stalk

We next asked, how the forming optic fissure margins, containing the pioneering edge cell population, are formed. Our data indicate, that next to the lens-averted domain of the optic vesicle, also the optic stalk contains cells of the future optic cup (Figure 3, Figure 4), in line with previous observations (Holt, 1980, Kwan et al., 2012). We recently found that TGFβ is important for optic fissure fusion (Knickmeyer et al., 2017 Biorxiv) and observed active TGFβ signaling within the optic fissure margins in zebrafish, using an *in vivo* signaling reporter (Knickmeyer et al., 2017, Heermann et al., 2014, Biorxiv). Here, we made use of this TGFβ reporter line to investigate whether the signaling domain is extending into the margins by TGFβ signaling activation in the margin cells, or by secondarily integration of cells, in which TGFβ signaling was activated, into the margins. Our data indicate that the latter is the case. Cells in which TGFβ signaling was activated are being brought into the fissure margins secondarily (Figure 5, supplemental movie files 9–11). Notably, these cells were stemming from the optic stalk (Figure 5 A, B, arrowhead), which is predominantly connected to the nasal ventral optic cup domain and the nasal fissure margin (Figure 5 A, B). However, the TGFβ signaling positive cells also largely integrated into the forming optic nerve head region (Figure 5 C, D) and via this region extended into the temporal fissure margin (Figure 5 C, D, H, arrow).

**Figure 5.**
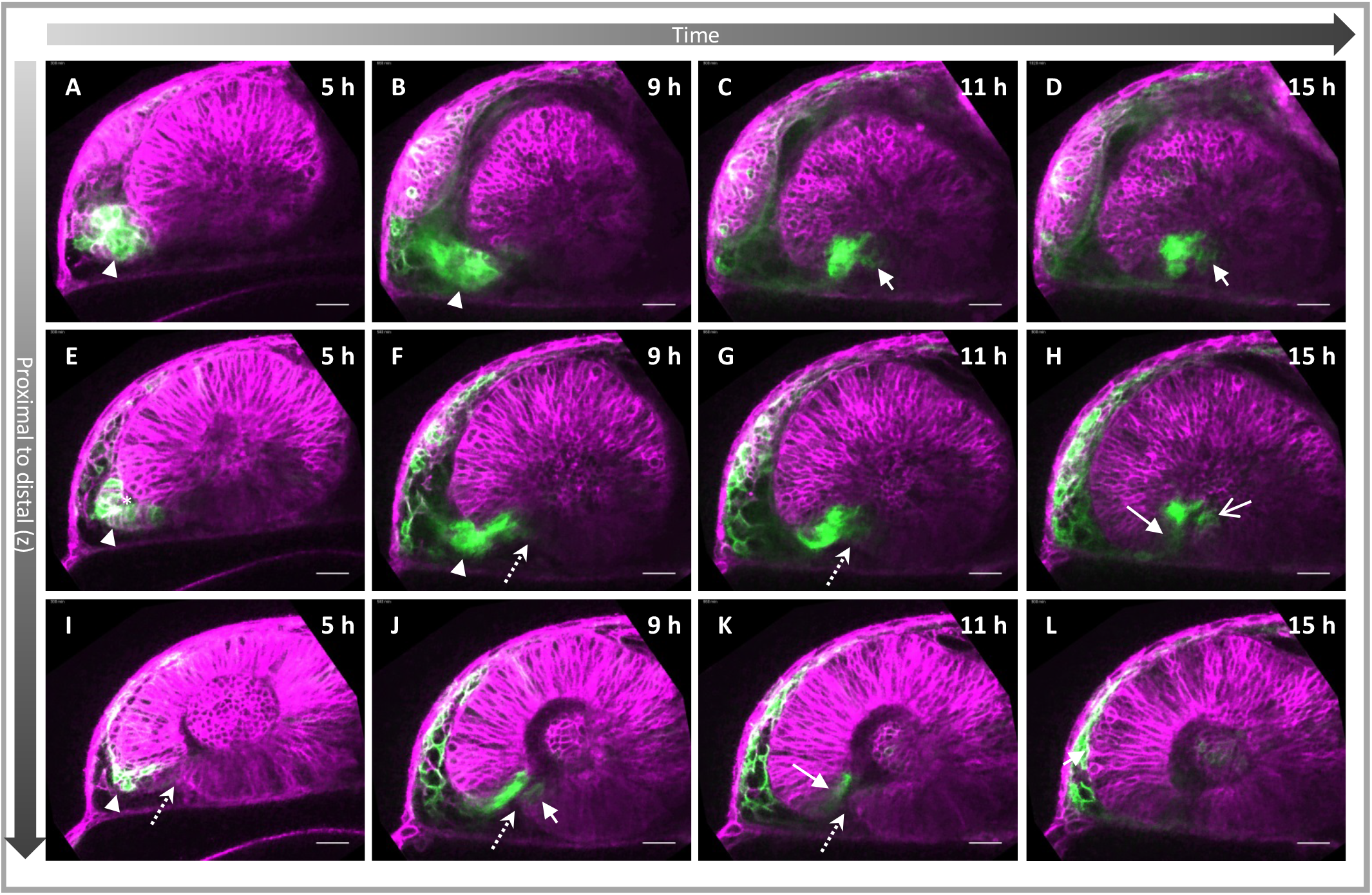
TGFβ signaling positive cells are secondarily added to the optic fissure margins. (A-L) 4D data set of a developing optic cup, TGFβ reporter (green), cell membranes (lyntdTomato, magenta), presented are 3 different optical section planes (top to bottom) over time (left to right). TGFβ reporter activity in the optic stalk (A, E, I, arrowhead), TGFβ signaling positive cells move from the optic stalk into the optic cup (C-K, arrow). The optic cup is predominantly connected to the nasal fissure margin (A, B, E, F). TGFβ signaling positive cells populate the future optic nerve head region, the most proximal domain of the optic fissure (C, D, F, G). From here these cells also populate the temporal fissure margin (C, D, F, G, H, J). The optic fissure is marked with a dotted arrow. Scalebar: 25μm

In distal fissure domains, the reporter signal is lost over time, due to either a continuous flow of these cells into other domains, or due to a reduction in signaling activity inside these cells (Figure 5 J - K). The optic vesicle initially is originating from the eye field, situated in the forebrain, the late prosencephalon (Rembold et al., 2006). In the subsequent order of events, during optic cup morphogenesis, distinct domains of the optic cup seem shielded from potentially inductive signals, emanating from the region of the surface ectoderm and developing lens (Heermann et al., 2015), likely resulting in the formation of the ciliary marginal zone (CMZ) in the distal periphery of the optic cup. Here we show yet another domain of the optic cup, the optic fissure margins, which is only secondarily integrated on top, or better “on bottom”, importantly via the optic stalk. This distinct origin makes the margins of the optic fissure special in comparison to large parts of the dorsal optic cup. Notably, cells of the optic fissure margins are differentiating only very late and prior to optic fissure fusion the forming CMZ and the optic fissure margins are a continuous sheet in the distal and ventral domain of the optic cup, the region in which the distal and ventral neuroretinal flow was occurring.

## Summary and Conclusion

In this study, we addressed the morphogenetic events of the optic fissure generation and the consecutive optic fissure fusion by 4D *in vivo* time-lapse analyses using zebrafish (*Danio rerio*) as a model. We identified a distinct population of cells situated in the border region in between the prospective neuroretina and RPE, which we termed pioneering edge cells. These cells establish the contact in between the nasal and temporal optic fissure margins in an EMT-like process, using a part of a hyaloid vessel as a scaffold to bridge the gap. Finally, we show that the optic fissure is a product of a bilateral flow from the optic stalk and lens-averted optic cup domain. In addition, cells are added to the ventral optic cup, specifically to the optic nerve head and optic fissure margins from the region of the optic stalk.

Overall, we show that optic fissure closure is a multistep process, in which all steps must be accomplished. If not, distinct defects of optic fissure closure are the result. The important steps are, the dynamic morphogenesis of the fissure by a bilateral tissue flow, proceeding in the setup of the fissure margins, by late secondary integration of cells from the optic stalk and peaking in an EMT-like epithelial margin disassembly, consecutive fusion and re-epithelialization. Based on our findings, we suggest, that if this multistep process is hampered during the early steps, it is resulting in extended coloboma, if the process is hampered during fusion, a mild coloboma is the result.

Our findings with respect to optic fissure fusion, essentially show an epithelial fusion, which can likely be generalized and applied also to other processes of tissue fusion, e.g. palatal shelf fusion. Furthermore, since EMT is also a hallmark of cancer formation and neural crest delamination, it is conceivable that these processes share morphological features with the EMT-like fissure margin disassembly during optic fissure fusion.

## Materials and Methods

### Zebrafish

**Husbandry**: Zebrafish (*Danio rerio*) were kept in closed stocks in accordance with local animal welfare law. The fish facility is under supervision of the local representative of the animal welfare agency. Fish were maintained in a constant recirculating system at 28°C on a 12h light/12h dark cycle. **Transgenic reporter zebrafish: TGFβ Reporter**: (*Tg*(*sbe:GFPcaax*) were generated previously, as described in Knickmeyer et al. (2017, Biorxiv, and in preparation), ***rx2* Reporter**: *tg(ola.rx2:egfp-caax)* were described and used previously (Heermann et al., 2015), **ubiquitous nuclear and membrane reporter**: The double transgenic line containing, *Tg*(*actb2:H2BGFP*/ *Tg*(*actb2:lyntdTomato*), greated by Burkhard Höckendorf in the lab of Jochen Wittbrotd and kept in a AB/Casper background. The line was kindly provided by the lab of Jochen Wittbrodt. **Transient labeling of Zebrafish**: Where indicated, mRNA for either H2BRFP (nuclear localized RFP) (kindly provided by the Lab of Lucia Poggi) (200 ng/μl), H2BGFP (nuclear localized GFP) (50ng/μl), lyntdTomato (membrane localized tandemTomato) (150ng/μl) was injected into 1–8 cell staged zebrafish embryos enabling 4D imaging of mosaic or ubiquitous labeled ZF. For mosaic labeling of POM cells a construct *T2*(*sox10:GFPcaax cmlc2:GFP*) was assembled in a Gateway reaction, using a Tol2 destination vector including *cmlc2:eGFP* (Kwan et al., 2007), a 5’Entry vector containing the *sox10* promoter (kindly provided by Bruce Apple), a middle entry vector containing GFPcaax (pENTR D-TOPO (ThermoFisher Scientific) and a 3’Entry vector with a polyadenylation site (Kwan et al., 2007). The construct (10ng/μl) was injected into wild type zebrafish zygotes together with Tol2 transposase mRNA (7ng/μl).

### Immunohistochemistry on sections

Immunohistochemistry was performed according to a standard protocol. Briefly, embryos at 30hpf were fixed in 4%PFA, washed, and transferred to 30% sucrose. Embryos were subsequently embedded (laterally) in Tissue Tec (Sigma) and sectioned (20μm) with a Cryostate CM3050s (Leica) at −25°C and dried overnight. Sections were blocked (BSA [1%], DMSO [1%], Triton X-100 [0.1%], NGS [4%], PBS [1×]). Embryos were incubated in primary antibody solution (anti Laminin Ab-1, 1:100, Neo Markers Fremont CA) in blocking solution overnight. Samples were washed and incubated in secondary antibody solution (anti rabbit Alexa 488 1:500, Invitrogen) with DAPI (stock: 2 μg/ml, 1: 500) for 2 hours. Consecutively, samples were washed and mounted in Moviol (Sigma) for microscopy.

## Microscopy

### Sections

Sections were imaged with a Leica TCS SPE confocal microscope with a z spacing of 4 μm 3 consecutive sections are depicted.

### Time-lapse imaging

Time-lapse imaging was performed with Leica TCS SP8 setups with 2 internal hybrid detectors, and a 40x long distance objective (water immersion), as immersion medium Immersol (Carl Zeiss) was used. For time-lapse imaging, embryos at appropriate stages were embedded in 1% low melting agarose in glass bottom dishes (MatTek, Ashland, MA) and covered with zebrafish medium, including tricaine for anesthesia. Confocal stacks were taken every 10min with a resolution of 1024 x 1024 and 3 μm z-spacing. Left and right eyes were used and oriented to fit the standard lateral view. For imaging of ZF older than 24 hours, fish were treated with PTU before imaging.

### Processing of time-lapse imaging data

The stacks were evaluated with the FIJI software (Schindelin et al.,2012). For denoising of the movies, the PureDenoise plugin (Luisier et al., 2010) was used with 4 Cycle-spins and 3 frames Multiframe.

### Single Plane Illumination (SPIM/ lightsheet) imaging

Time-lapse spim imaging was performed with a Leica TCS SP8 setup that was upgraded with a DLS (Digital lightsheet). A 5x illumination objective, a 25x dipping lens together with 2.5mm mirror caps were used to obtain a lightsheet. A stack was taken every 5 min. For imaging, glass bottom dishes (MatTek, Ashland, MA) were coated with 2% Agarose. Embryos were embedded in 1% low melting agarose and placed on the coating. With razorblades 2 notches were cut in the agarose, that the embryo was placed on a 2mm wide strip that could fit between the mirror caps. The embryo was covered with zebrafish medium, including tricaine for anesthesia. A stack was taken every 5 min. Left and right eyes were used and oriented to fit the standard lateral view.

### Tracking and visualization

Tracking of cells was performed with MTrackJ (Meijering et al., 2012). Tracks were visualized using custom made ImageJ plugins as in (Heermann et al., 2015). The overlay images of tracks and raw data were used to produce maximum intensity projections for Figure 5 and supplemental movie file 8.

### Quantitative analysis contact height between the fissure margins

To analyze the height of the contact between the fissure margins, in 7 embryos one distal optical section was used to measure the distance between to circles, one circle was placed to mark the lens-averted border of the optic fissure, another was placed inside the optic cup to mark the lens-facing border of the fissure. The complete distance was measured and set as 100%. The pioneering edge cell was virtually split in half and the distance to the respective borders was measured from that point.

## Acknowledgements

We want to thank Bruce Appel for generously providing the sox10 cis regulatory element containing p5E vector, Burkhard Höckendorf and the Wittbrodt Lab for providing transgenic lines and Lucia Poggi for sharing mRNAs.

## Funding

LS was funded by a fellowship of the Hartmut-Hoffman-Berling International Graduate School (HBIGS). This project was funded by the “Forschungskommission” of the University Freiburg (HEE1079/16).

## Figure Legends

**Figure S1.**
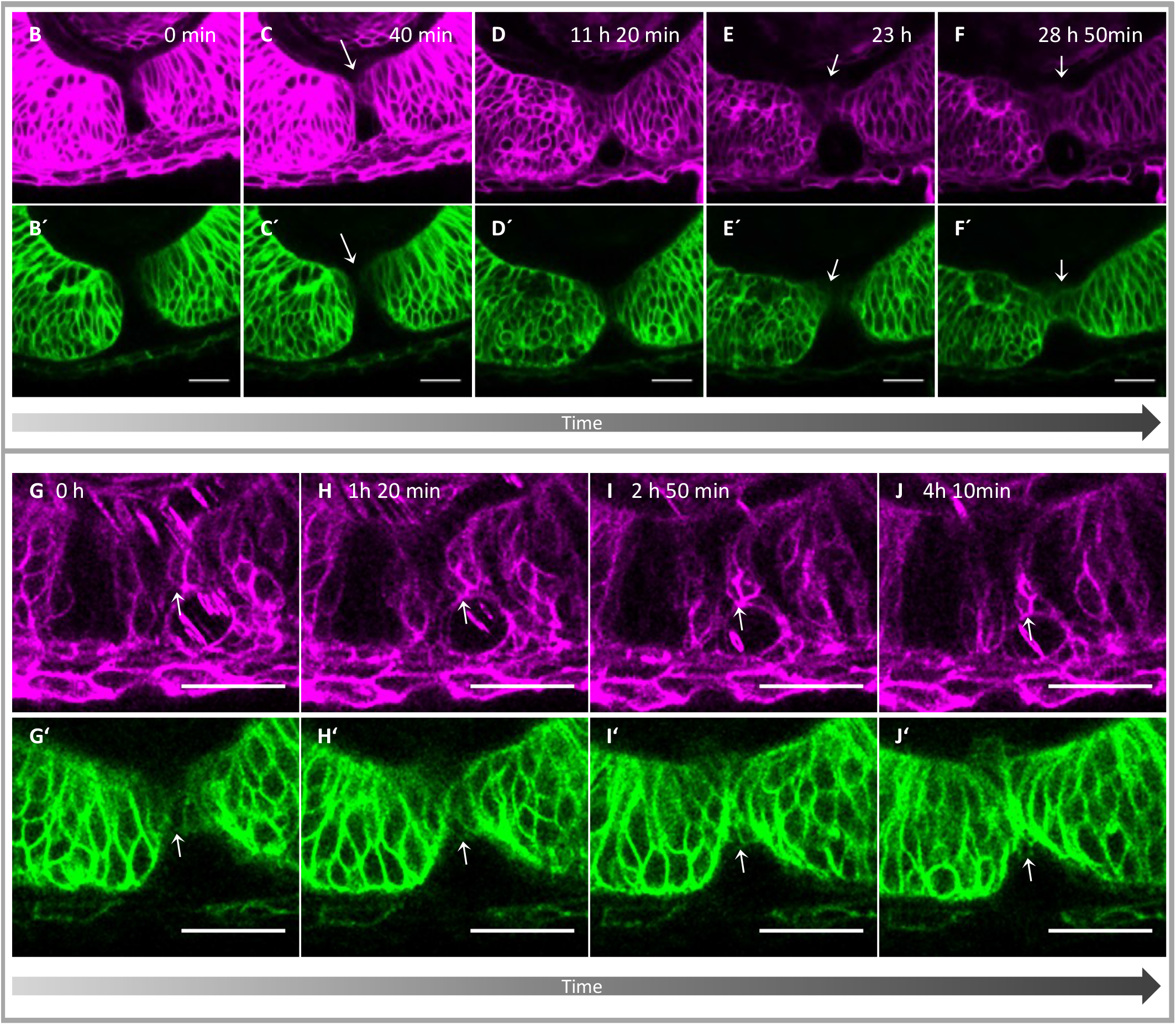
(A-J) single channels of the respective merged pictures form Figure 1. Please note the mosaic labeling with lyntdTomato (F-I).

**Figure S3.**
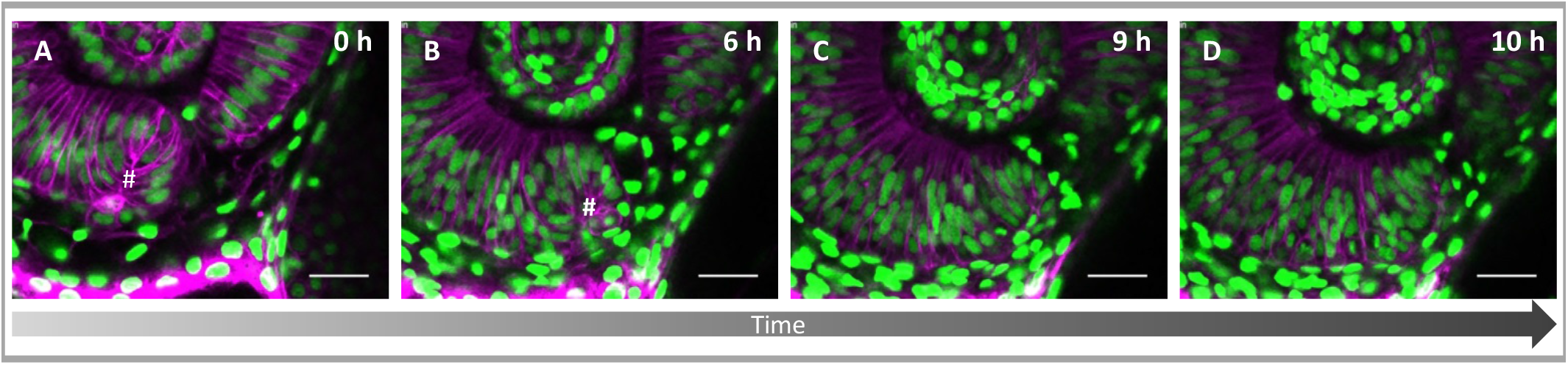
A vortex of cells in the nasal ventral optic cup. (A-D) A cellular vortex (A,B, #) is visible in the nasal optic fissure margin, where cells from the RPE domain are moving from the lens-averted layer into the lens-facing layer through the optic fissure and the distal rim. Scalebar: 25μm

## Movie Legends

**Supplemental Fig 1 Movie 1**: related to Figure 1 B-F **Pioneering edge cell mediates contact between the fissure margins and reactivates rx2 expression**, visualized by lyntdTomato mRNA into *Tg*(*rx2:GFPcaax*) (orientation as in Figure 1A) imaging starts at 32 hpf framerate 1/10 min).

**Supplemental Fig 1 Movie 2**: related to Figure 1 G-J **Pioneering edge cell progeny integrates into RPE**, visualized by lyntdTomato mRNA into *Tg*(*rx2:GFPcaax*) (orientation as in Figure 1A) imaging started at 32 hpf the movie starts 58 hpf framerate 1/10 min).

**Supplemental Fig 2 Movie 3**: related to Figure 2 (A-C) **Optic Fissure is closing over the hyaloid vessel** visualized by lyntdTomato mRNA into *Tg*(*fli:eGFP*) (orientation as in Figure 1A) imaging starts at around 35 hpf framerate 1/10 min).

**Supplemental Fig 2 Movie 4**: related to Figure 2 **POM cell is moving around the optic cup and into the optic fissure** visualized by H2BRFP mRNA, Tol2 mRNA and a Plasmid containing the *Tol2-Sox10:GFPcaax*, imaging starts at 24hpf, framerate 1/10 min).

**Supplemental Fig 3 Movie 5** related to Figure 4 A-C **Proximal Optic Fissure development**. visualized *Tg*(*bact:H2BGFP*) and *Tg*(*bact:lyntdTomato*), imaging starts at 18hpf, framerate 1/10 min and imaged at 28°C over night.

**Supplemental Fig 3 Movie 6** related to Figure 4 D-F Distal **Optic Fissure development**. visualized *Tg*(*bact:H2BGFP*) and *Tg*(*bact:lyntdTomato*), imaging starts at 18hpf, framerate 1/10 min and imaged at 28°C over night.

**Supplemental Fig 3 Sup Movie 7**: related to Figure S 4 **Nasal Vortex. Cells flow from lens-averted layer in lens-facing layer** visualized by *Tg*(*bact:H2BGFP*) and *Tg*(*bact:lyntdTomato*), imaging starts at around 26 hpf, framerate 1/10 min and imaged at 28°C over night.

**Supplemental Fig 4 Movie 8**: related to Figure 5 (E-H) **single cells moving around the ventral temporal rim**, visualized by H2BRFP mRNA in mosaic/sparse labeling, imaging starts at 16,5hpf, framerate 1/5 min and imaged at 25°C over night. Max projection of 70 optical sections corresponding to 70μm (z-spacing was 1μm)

**Supplemental Movie 9 -11**: related to Figure 5 (K-V) **TGFβ signaling positive cells are integrated into the optic fissure margins (9–11, proximal to distal)**, visualized by lyntdTomato mRNA into *Tg*(*SBE:GFPcaax*), imaging starts at 18hpf, framerate 1/10 min.Every movie represents 1 optical section between the optical sections 21 μm distance

